# Genetic parameters and genome-wide association analysis of service sire effect on litter size and its relationship with boar semen quality in three terminal sire lines

**DOI:** 10.1101/2025.08.01.668164

**Authors:** Ching-Yi Chen, Daniela Lourenco, Michael Kleve-Feld, Adria S. Bhatnagar, Justin W. Holl

**Affiliations:** The Pig Improvement Company, Genus plc, Hendersonville, TN, United States; Department of Animal and Dairy Science, University of Georgia, Athens, GA, United States

## Abstract

This study aimed to estimate the genetic parameters for service sire effects on the number born alive (NBA) and its relationship with semen quality traits. Data of 6,416, 23,188, and 48,890 litter size records collected between 2020 and 2024 from three purebred terminal sire lines were analyzed. The number of sows was 3,071, 11,819, and 24,089, with 197, 554, and 891 service sires for the three respective lines. There were 1,424,858, 4,344,630, and 2,146,583 animals in the pedigree of which 67,990, 259,250, and 365,392 were genotyped and imputed up to 50K SNPs. The service sire and dam effects were modeled as additive genetic effects, considering a covariance structure among them, given by the relationship matrix. The model also included fixed effects of contemporary group and parity group with random permanent environmental effects for service sire and dam. Genetic parameters were estimated using the AIREML option in the BLUPF90+ software, and GEBVs were generated by ssGBLUP with the algorithm for proven and young (APY). Heritability for the service sire effect ranged from 0.01 to 0.03, and the heritability for the dam effect ranged from 0.09 to 0.15, with genetic correlations ranging from -0.20 to 0.37. A single-step genome-wide association study (ssGWAS) was also performed, and no strong signals were detected for service sire effects on NBA. Sperm motility (MOT) and total abnormal morphology (ABN_MOR) GEBVs from the three lines were estimated based on about 57,500 to 608,996 ejaculates recorded using the computer-assisted semen analysis (CASA) system from 1,698 to 29,095 boars collected from 2013 to 2025. Heritability estimates ranged from 0.10 to 0.12 for motility and from 0.17 to 0.28 for total abnormal morphology. The correlations between service sire GEBV for NBA and semen quality were low but in the favorable direction. Results suggest that although paternal genetic contributions to litter size were small compared to maternal genetic contributions, selecting on service sire effects on litter size in addition to semen quality traits will improve overall reproductive success.

## Background

Reproductive performance in terminal sire lines is typically focused on boar fertility traits, such as semen quality, whereas traits like litter size are primarily addressed through maternal line breeding objectives. However, to improve overall male reproductive success and better capture a terminal boar’s contribution to downstream reproductive outcomes, the paternal genetic influence on litter size is of particular interest, especially in artificial insemination systems, where a single sire can impact a large number of litters. Although litter size is largely determined by maternal components, service sires can also influence it through biological and genetic aspects [1–3]. Recognizing and integrating these paternal contributions can enhance reproductive efficiency and support the overall success of pig breeding programs.

Previous studies have investigated the impact of the service sire effect on litter size in swine, either by modeling it as a random non-genetic effect uncorrelated with sows [4–8] or by modeling it as additive genetic effect correlated with sows [9,10]. With the accessibility of Computer-Assisted Sperm Analysis (CASA) technology, artificial insemination centers can routinely assess boar semen quality through automated estimates of various parameters, including sperm motility and morphology. Literature on the genetic parameters of semen quality traits revealed low to moderate heritability [7,11–13] and a wide range of genetic correlations with sow litter size traits [7,12,14].

There are limited reports on investigating the service sire effects on litter size incorporated with genomics information, including single-step genomic best linear unbiased prediction (ssGBLUP) [15] and single-step approach genome-wide association study (ssGWAS) for large genotyped swine populations.

The ssGWAS was developed by Wang et al. [16] to estimate marker effects from ssGBLUP. To measure statistical significance for marker effect, later Aguilar et al. [17] expanded the method proposed by Lu et al. [18] to derive p-values for single-marker GWAS within the ssGWAS framework. The algorithm was validated using a large dataset from American Angus population. All p-values were obtained in a single run with negligible computational time. Misztal et al. [19] indicated that there was a soft limit of about 100K genotyped animals when obtaining the p-value by inverting the left-hand side of the mixed model equations. An alternative approach to bypass the inverse of the left-hand side components was developed by Bermann et al. [20] to eliminate the bottleneck in approximating reliabilities for large-scale genotyped populations. In their study, they extended the work from Misztal and Wiggans [21] on approximating prediction error (co)variances of GEBVs for the APY core set. Their work has been validated with about 330K genotyped animals from American Angus population. Leite et al. [22] applied the approximation of p-value for ssGWAS on 450K genotyped animals.

In this study, we aim to evaluate the paternal influence on litter size and its genetic relationship with semen quality traits in terminal sire lines and to investigate significant genomic regions associated with service sire effects on litter size using ssGWAS. To our knowledge, our study is the first ssGWAS to investigate the service-sire effect on litter size in large genotyped swine populations.

## Methods

### Data

Data was provided by PIC (a Genus company, Hendersonville TN, USA). For litter size analysis, the data set contained 6,416, 23,188, and 48,890 farrowing records collected between 2020 and 2024. There were 3,071, 11,819, and 24,089 sows sired by 197, 554, and 891 boars from terminal sire lines A, B, and C, respectively. For semen quality analysis, the data set contained 57,500, 608,996, and 346,059 ejaculates recorded collected between 2013 and 2025. There were 1,698, 29,095, and 18,726 boars from terminal sire lines A, B, and C, respectively. Semen quality of motility and total abnormal morphology were measured using the computer-assisted semen analysis (CASA) system. Data quality controls were applied to exclude motility < 50% and abnormal morphology >40%. The total abnormal morphology were calculated based on tail miscellaneous abnormalities, head abnormalities, distal cytoplasmic droplets, and proximal cytoplasmic droplets.

The pedigree included 1,424,858, 4,344,630, and 2,146,583 animals for lines A, B, and C, respectively, among which 67,990, 259,250, and 365,392 animals were genotyped and imputed up to 50K SNPs. Number of animals used in the analysis and the descriptive statistics for the 3 lines are shown on Table 1. The average number of piglets born alive ranged from 8.57 to 9.38. Average sperm motility ranged from 89.22% to 90.91%, while the average total abnormal morphology rate ranged from 9.52% to 12.17%.

**Table 1.** Descriptive statistics.

| Line | N | Mean | SD | Min | Max |
| --- | --- | --- | --- | --- | --- |
| NBA, pig |  |  |  |  |  |
| A | 6,416 | 8.57 | 3.24 | 0 | 18 |
| B | 23,188 | 9.04 | 2.83 | 0 | 20 |
| C | 48,890 | 9.38 | 3.00 | 0 | 22 |
| MOT, % |  |  |  |  |  |
| A | 57,500 | 89.97 | 6.43 | 50 | 100 |
| B | 608,996 | 90.91 | 5.56 | 50 | 100 |
| C | 346,059 | 89.22 | 6.51 | 50 | 100 |
| ABN_MOR, % |  |  |  |  |  |
| A | 57,500 | 11.25 | 6.51 | 0 | 40 |
| B | 608,996 | 9.52 | 6.20 | 0 | 40 |
| C | 346,059 | 12.17 | 8.06 | 0 | 40 |

### Models and statistical analyses

Genetic parameters were estimated using BLUPF90+ with AIREML option [23].

### Litter Size Genetic Parameters

Analyses were performed using a single-trait model:

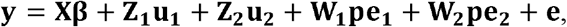

Where **y** is the vector of phenotypes for NBA; **β** is the vector of fixed effects of parity group and contemporary group defined as farm and farrowing year and month; **u**_**1**_ and **u**_**2**_ are the vectors of random additive service sire and dam genetic effects; **pe**_**1**_ and **pe**_**2**_ are the vectors of random service sire and dam permanent environmental effects; **e** is the vector of random residual effects; **X, Z**_**1**_, **Z**_**2**_, **W**_**1**_, and **W2** are the incidence matrices relating records in vector **y** to effects in **β, u**_**1**_, **u**_**2**_, **pe**_**1**_, and **pe**_**2**_ respectively. Random effects were assumed to follow a multivariate normal distribution with a mean of zero and covariance structure as outlined below:

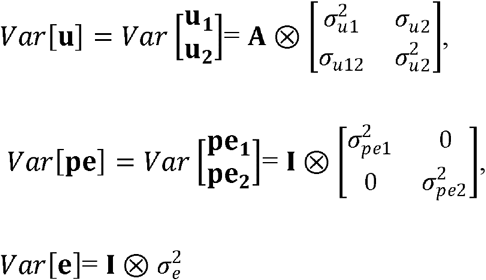

where **A** is the numerator relationship matrix; **I** is the identity matrix; 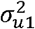 and 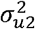 are the additive genetic variances of service sire and dam with *σ*_u12_ as the genetic covariance; 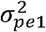 and 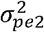 are the permanent environmental variances of service sire and dam; 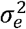 is the residual variance.

### Semen Quality Genetic Parameters

Analyses were performed using a two-trait model:

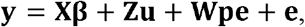

where **y** is the vector of phenotypes for MOT and ABN_MOR; **β** is the vector of fixed effects of contemporary group defined as farm and year and month at collection and the linear covariates of boar age at collection and rest days between two collections; **u** is the vector of random additive genetic effects; **pe** is the vector of permanent environmental effects; **e** is the vector of random residual effects; **X, Z**, and **W** are the incidence matrices relating semen records in vector **y** to effects in **β, u**, and **pe**, respectively. Random effects were assumed to follow a multivariate normal distribution with a mean of zero and covariance structure as outlined below:

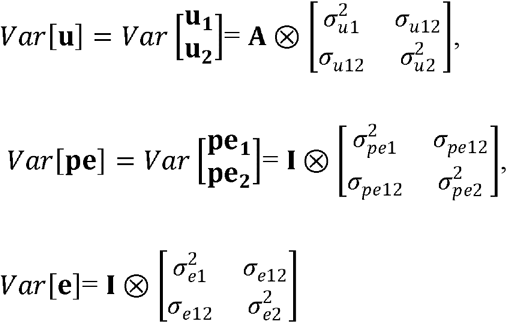

where **A** is the pedigree-based numerator relationship matrix; **I** is the identity matrix; ;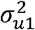 and 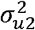 are the additive genetic variances for MOT and ABN_MOR with *σ*_*u*12_ as the genetic covariance; 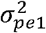 and 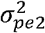 are the permanent environmental variances with covariance of 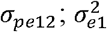 and 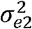 are the residual variances with covariance of *σ*_*e*12_.

### Breeding Value Estimation

Genomic estimated breeding values (GEBVs) were generated using ssGBLUP, which combines the pedigree-based relationship matrix, **A**, and the genomic relationship matrix (**G**) into a unified relationship matrix (**H**). Genomic relationship matrices compatible with pedigree information have been investigated and developed [15,24–28]. The inverse matrix, **H**^-1^, described as in Aguilar et al. [15] and Christensen and Lund [29] is:

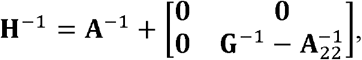

Where **A**^-1^ is the inverse of the pedigree relationship matrix, 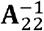 is the inverse of the pedigree relationship matrix for genotyped animals, and **G**^-1^ is the inverse of the genomic relationship matrix. Lourenco et al. [23] reviewed that the implementation of algorithm for proven and young (APY) in ssGBLUP enables highly efficient GEBV computation for large numbers of genotyped animals by constructing a sparse representation of **G**^-1^ and using an efficient algorithm to compute 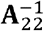 [30–32]

Pocrnic et al. [33] proposed the number of core animals could be defined according to the dimensionality of the genomic information. For pigs, the optimal number of core animals was reported as 6,000, which was defined based on the number of eigenvalues explaining 98% of the variation in **G**. In this study, the core animal set consisted of service sires along with randomly selected animals, with a combined total of 20,000 individuals.

### Genome-Wide Association Study

With large number of genotyped animals being used in the study, we conducted ssGWAS using method proposed by Leite et al. [22] to approximate marker p-values to avoid the expensive computational cost on inverting high dimensional dense matrices associated with genotyped animals. Combining with APY approach, only relationships and estimated breeding values for core animals are needed for estimating marker effects. The obtained p-values were adjusted using the Bonferroni correction for multiple testing, with a significance threshold set at the 5% genome-wide level [17,22]

## Results

### Genetic parameters

Estimates of variance components, heritability, and genetic correlation for NBA are presented in Table 2. For line A, the full model including permanent environmental effects for both service sire and sow failed to converge. Convergence issues remained when the model retained the service sire’s permanent environmental effect but excluded that of the sow. In contrast, convergence was achieved when the service sire’s permanent environmental effect was excluded and only the sow’s was retained. Therefore, a reduced model excluding the service sire’s permanent environmental effect was used for line A, while the full model was applied for lines B and C. The genetic variances attributed to the service sire were relatively small, accounting for approximately 27%, 7%, and 19% of the sow genetic variances for lines A, B, and C, respectively. The genetic covariance between service sire and sow for line A was negative but with high SE (−0.11±0.13). Positive genetic covariances were observed for line B and C with small SE (0.11±0.03 and 0.08±0.04). The permanent environment variances for service sire were less than 10% of those observed for the sow. The heritability estimate for the service sire were 0.03, 0.01, and 0.02 for lines A, B, and C, respectively. Corresponding heritability estimates for the sow were 0.11, 0.15, and 0.09. The estimates of genetic correlations between service sire and sow effects were -0.20, 0.37, and 0.23 for lines A, B, and C, respectively. Repeatability for service sire effects were near zero for all lines whereas the repeatability for sow effects were ranged from 0.18 to 0.24.

**Table 2.** Estimates of genetic parameters (SE) for NBA.

|  | A | B | C |
| --- | --- | --- | --- |
| $\sigma_{u1}^2$ | 0.28 (0.08) | 0.08 (0.03) | 0.15 (0.04) |
| $\sigma_{u12}$ | -0.11 (0.13) | 0.11 (0.04) | 0.08 (0.04) |
| $\sigma_{u2}^2$ | 1.04 (0.22) | 1.16 (0.11) | 0.79 (0.07) |
| $\sigma_{pe1}^2$ | NA | 0.01 (0.02) | 0.06 (0.04) |
| $\sigma_{pe2}^2$ | 0.81 (0.19) | 0.76 (0.09) | 0.83 (0.07) |
| $\sigma_e^2$ | 7.32 (0.18) | 5.95 (0.08) | 6.92 (0.06) |
| $h_1^2$ | 0.03 | 0.01 | 0.02 |
| $h_2^2$ | 0.11 | 0.15 | 0.09 |
| $rep_1$ | NA | 0.01 | 0.02 |
| $rep_2$ | 0.20 | 0.24 | 0.18 |
| $r_g$ | -0.20 | 0.37 | 0.23 |

$\sigma_{u1}^2, \sigma_{u2}^2, \sigma_{pe1}^2, \sigma_{pe2}^2, \sigma_e^2$ : service sire additive genetic, dam additive genetic, service sire permanent environment, dam permanent environment, and residual variances; $h_1^2, h_2^2$ : heritability for service sire and dam; $rep_1, rep_2$ : repeatability for service sire and dam; $\sigma_{u12}, r_g$ : genetic covariance and correlation between service sire and dam. Standard errors in parentheses.

Estimates of variance components, heritability, and genetic correlation for MOT and ABN_MOR are presented in Table 3. For MOT, estimates of genetic variances were similar across the three lines (3.24 to 4.17) with larger phenotypic variance of 42.92 for line A and smaller phenotypic variances of 26.79 and 31.39 for lines B and C. For ABN_MOR, the estimate of genetic variances was nearly doubled in line C (14.13) compared to other two lines (7.76 and 8.08) with similar phenotypic variances for lines A and C (45.30 and 51.01) and smallest phenotypic variances of 32.94 for line B. Overall, line B has the smaller non-genetic variances among the three lines. The heritability estimates for MOT were 0.10, 0.12, and 0.11 for lines A, B, and C, respectively. For ABN_MOR, the heritability estimates were 0.17, 0.25, and 0.28 for the same lines. The genetic correlations between MOT and ABN_MOR were −0.34, −0.39, and −0.64 for lines A, B, and C, respectively. Estimates of repeatability ranged from 0.33 to 0.43 for MOT and from 0.50 to 0.54 for ABN_MOR. The GEBV correlations between service sire ranged from 0.06 to 0.13 for NBA and MOT, and from -0.08 to -0.12 for NBA and ABN_MOR. Those estimates were low but in the favorable direction (Table 4).

**Table 3.**
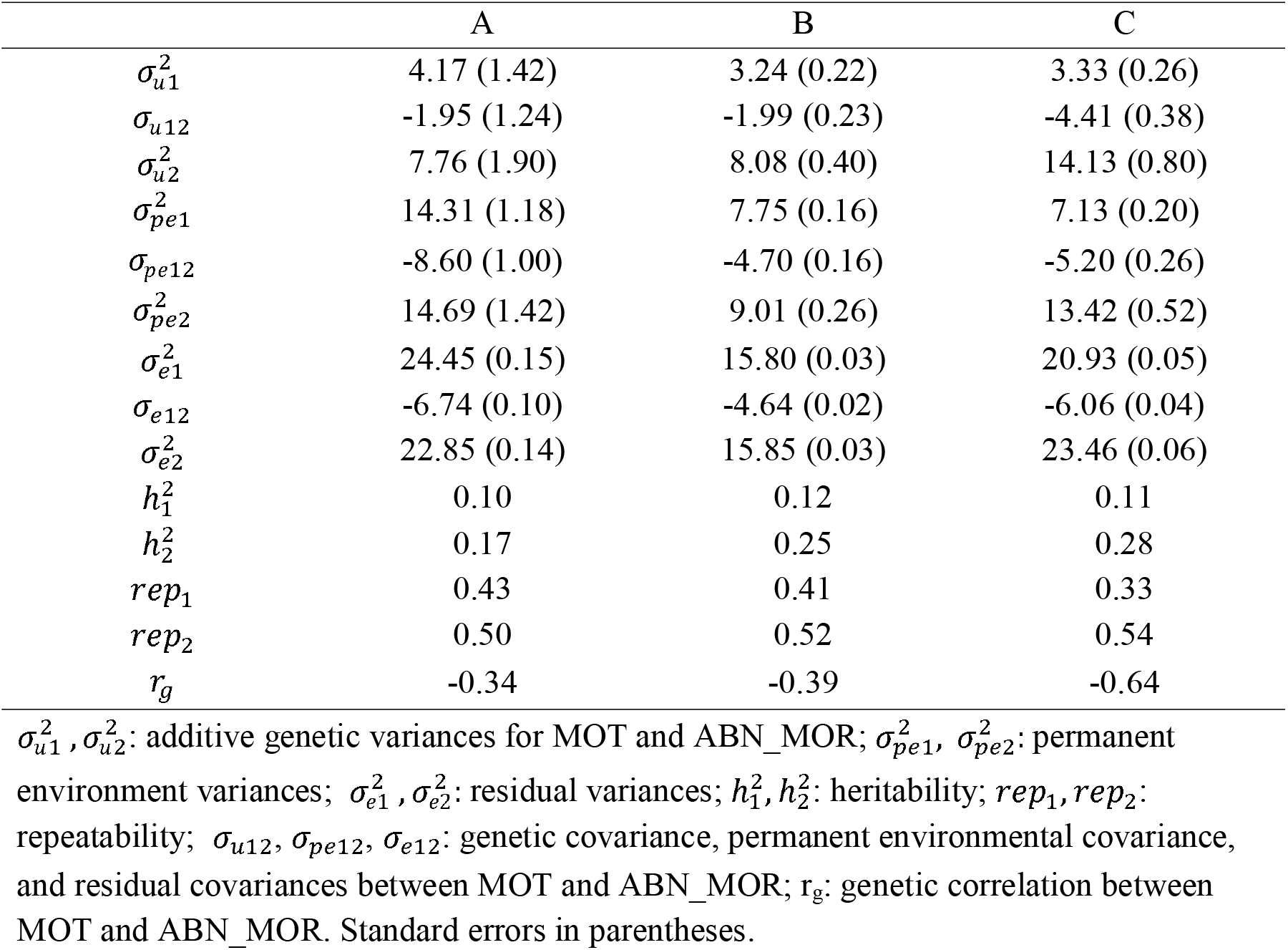
Estimates of genetic parameters (SE) for MOT and ABN_MOR.

|  | A | B | C |
| --- | --- | --- | --- |
| $\sigma_{u1}^2$ | 4.17 (1.42) | 3.24 (0.22) | 3.33 (0.26) |
| $\sigma_{u12}$ | -1.95 (1.24) | -1.99 (0.23) | -4.41 (0.38) |
| $\sigma_{u2}^2$ | 7.76 (1.90) | 8.08 (0.40) | 14.13 (0.80) |
| $\sigma_{pe1}^2$ | 14.31 (1.18) | 7.75 (0.16) | 7.13 (0.20) |
| $\sigma_{pe12}$ | -8.60 (1.00) | -4.70 (0.16) | -5.20 (0.26) |
| $\sigma_{pe2}^2$ | 14.69 (1.42) | 9.01 (0.26) | 13.42 (0.52) |
| $\sigma_{e1}^2$ | 24.45 (0.15) | 15.80 (0.03) | 20.93 (0.05) |
| $\sigma_{e12}$ | -6.74 (0.10) | -4.64 (0.02) | -6.06 (0.04) |
| $\sigma_{e2}^2$ | 22.85 (0.14) | 15.85 (0.03) | 23.46 (0.06) |
| $h_1^2$ | 0.10 | 0.12 | 0.11 |
| $h_2^2$ | 0.17 | 0.25 | 0.28 |
| $rep_1$ | 0.43 | 0.41 | 0.33 |
| $rep_2$ | 0.50 | 0.52 | 0.54 |
| $r_g$ | -0.34 | -0.39 | -0.64 |

$\sigma_{u1}^2, \sigma_{u2}^2$ : additive genetic variances for MOT and ABN\_MOR; $\sigma_{pe1}^2, \sigma_{pe2}^2$ : permanent environment variances; $\sigma_{e1}^2, \sigma_{e2}^2$ : residual variances; $h_1^2, h_2^2$ : heritability; $rep_1, rep_2$ : repeatability; $\sigma_{u12}, \sigma_{pe12}, \sigma_{e12}$ : genetic covariance, permanent environmental covariance, and residual covariances between MOT and ABN\_MOR; $r_g$ : genetic correlation between MOT and ABN\_MOR. Standard errors in parentheses.

**Table 4.** GEBVs correlation between service sire effect in NBA and boar semen traits.

| Line | N of boars | MOT | ABN_MOR |
| --- | --- | --- | --- |
| A | 197 | 0.13 | -0.12 |
| B | 554 | 0.06 | -0.08 |
| C | 891 | 0.10 | -0.12 |

### Genome-Wide Association Study

The Bonferroni-corrected rejection threshold for multiple testing at a nominal α □ = □ 0.05 was 5.90, 5.94, and 5.88 for lines A, B, and C, respectively. These thresholds were calculated as - log10(0.05/number of SNPs), based on 39,951, 43,505, and 38,228 SNPs for lines A, B, and C, respectively. Manhattan plots for genetic effect of service sire on NBA for the three lines are shown in Figure 1. No significant peaks were identified for lines A and C. There was one marker located on chromosome 12 at position near 48.29Mb having -log10(P-Value) =6.02 for the service sire genetic effect for line B. The Quantile–quantile plots of the genome-wide association studies for genetic effect of service sire on NBA are shown in Figure 2. The genomic inflation factor (λ) of the QQ plot regression were 0.842, 1.236, and 0.947 for line A, B, and C, respectively, indicating slight deflation for line A and slight inflation for line B.

**FIGURE 1.**
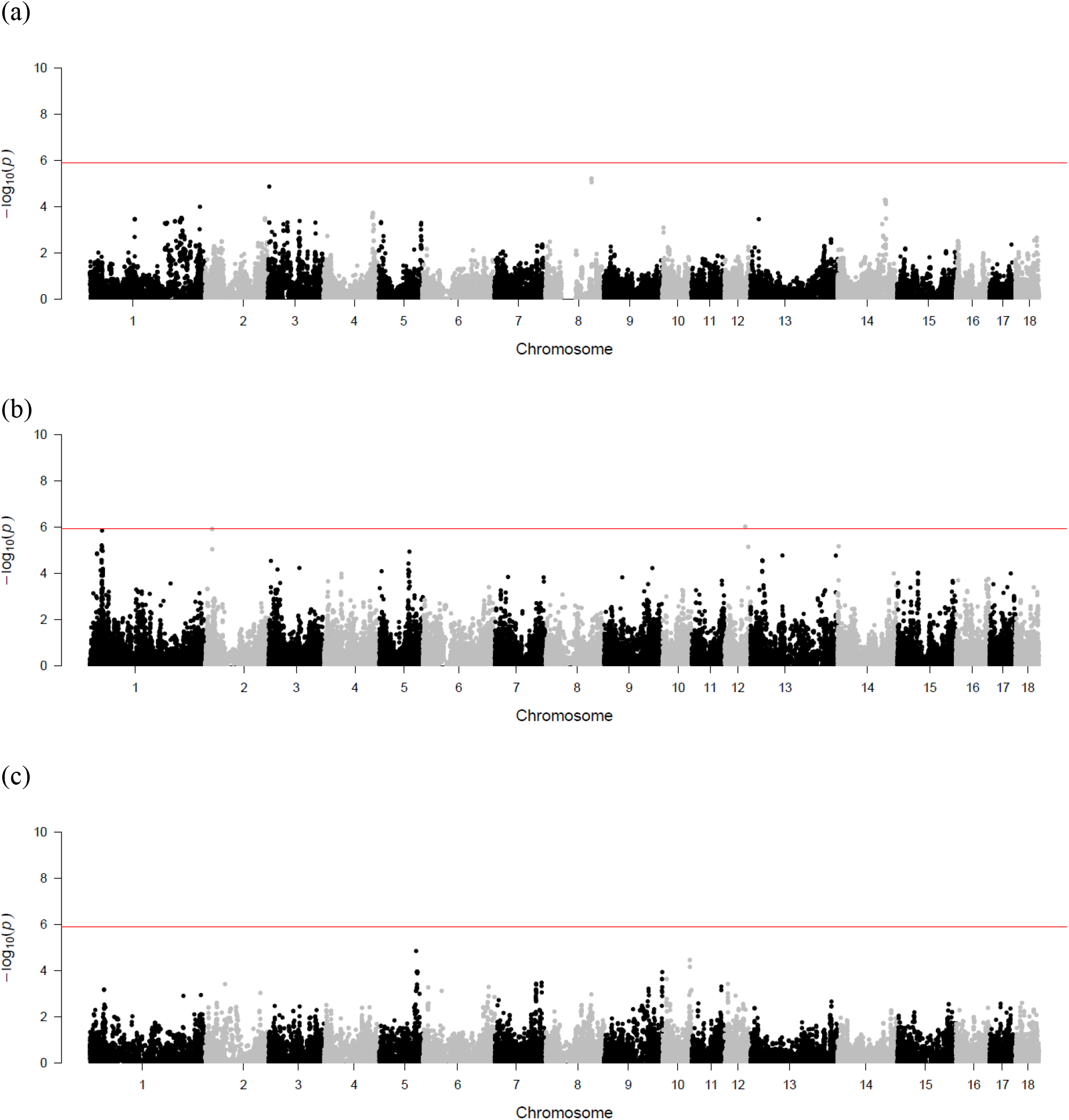
Manhattan plots of the single-step genome-wide association study for genetic effect of service sire on NBA for (a) Line A (b) Line B (c) Line C. The x-axis represents chromosome numbers with the genomic coordinates of SNPs along the axis. The y-axis represents the observed - log10 P-value. The red line indicates the Bonferroni-corrected significance threshold at a nominal α □ = □ 0.05, which was -5.90, -5.94, and -5.88 for lines A, B, and C, respectively.

**FIGURE 2.**
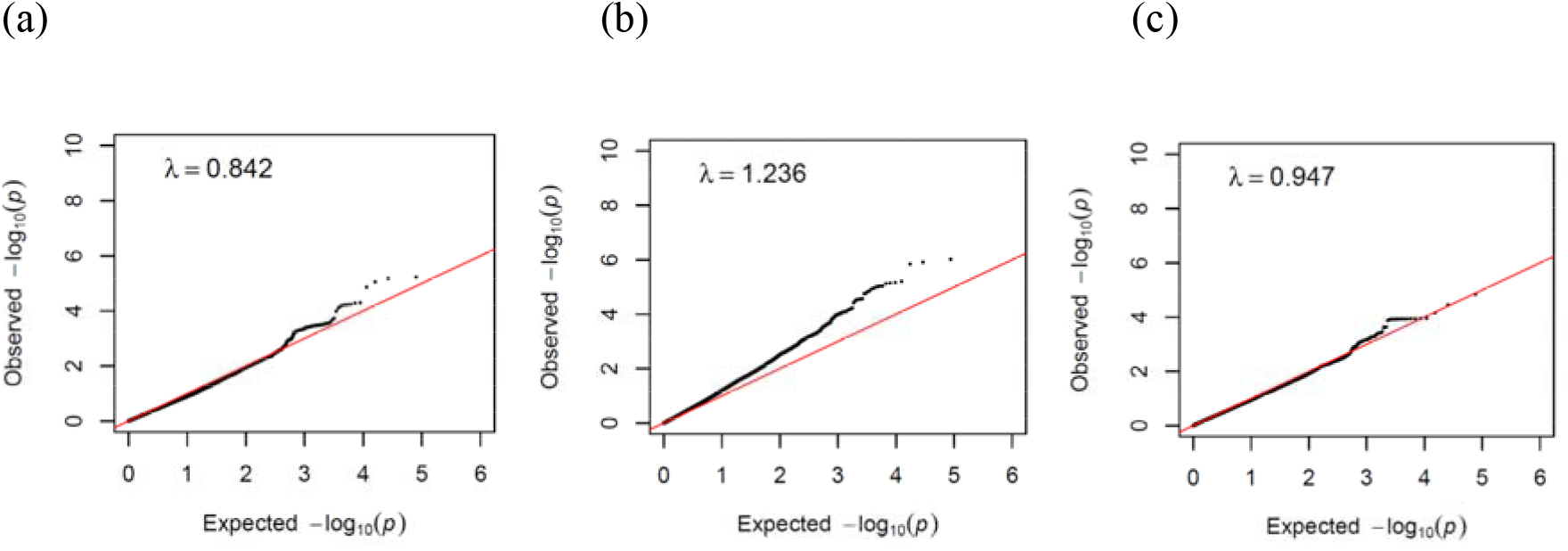
Quantile–quantile plots of the genome-wide association studies for genetic effect of service sire on NBA for (a) Line A (b) Line B (c) Line C.

**FIGURE 3.**
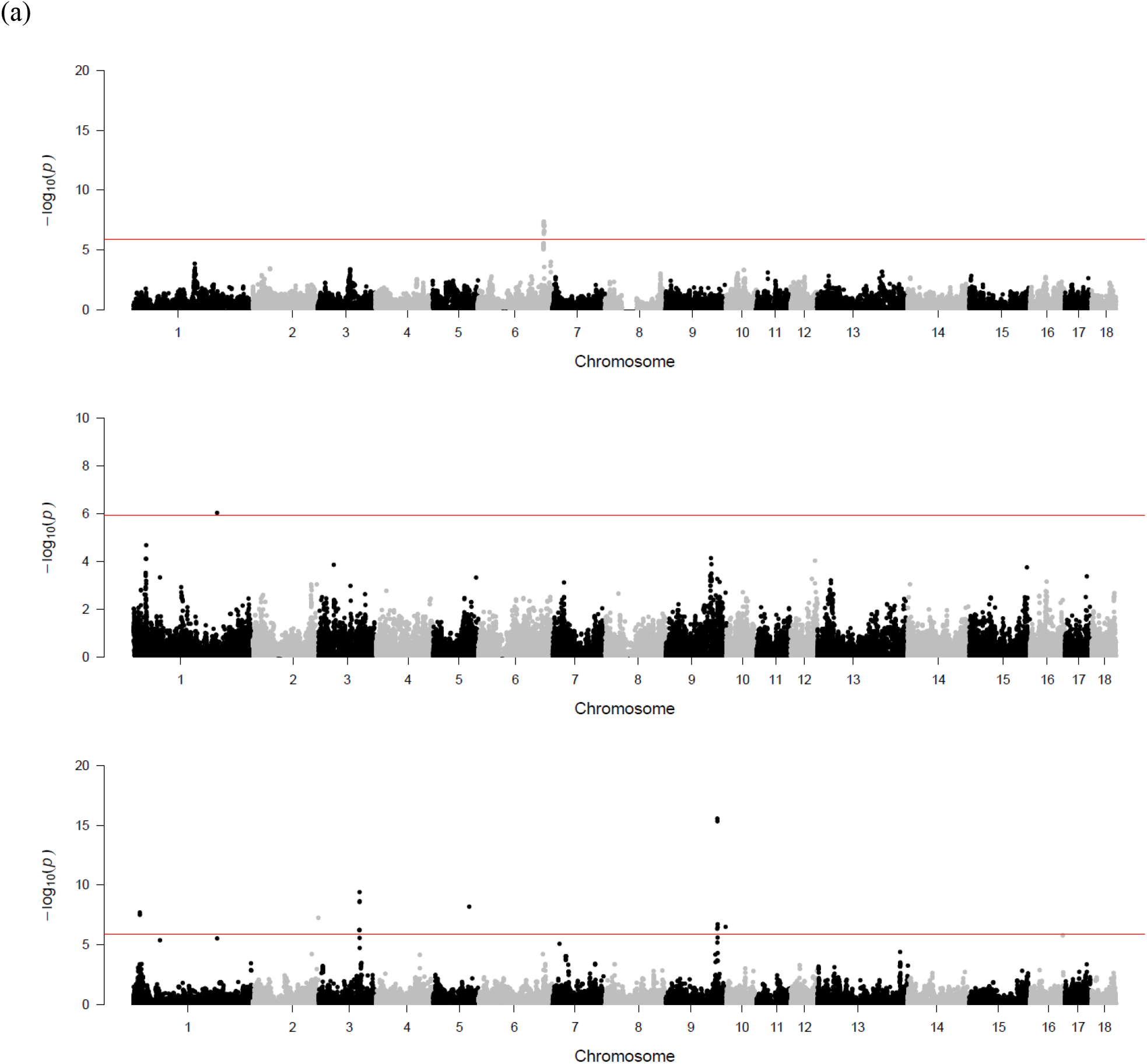
Manhattan plots of the single-step genome-wide association study for genetic effect of sow on NBA for (a) Line A (b) Line B (c) Line C. The x-axis represents chromosome numbers with the genomic coordinates of SNPs along the axis. The y-axis represents the observed -log10 P-value. The red line indicates the Bonferroni-corrected significance threshold at a nominal α □ = □ 0.05, which was -5.90, -5.94, and -5.88 for lines A, B, and C, respectively.

**FIGURE 4.**
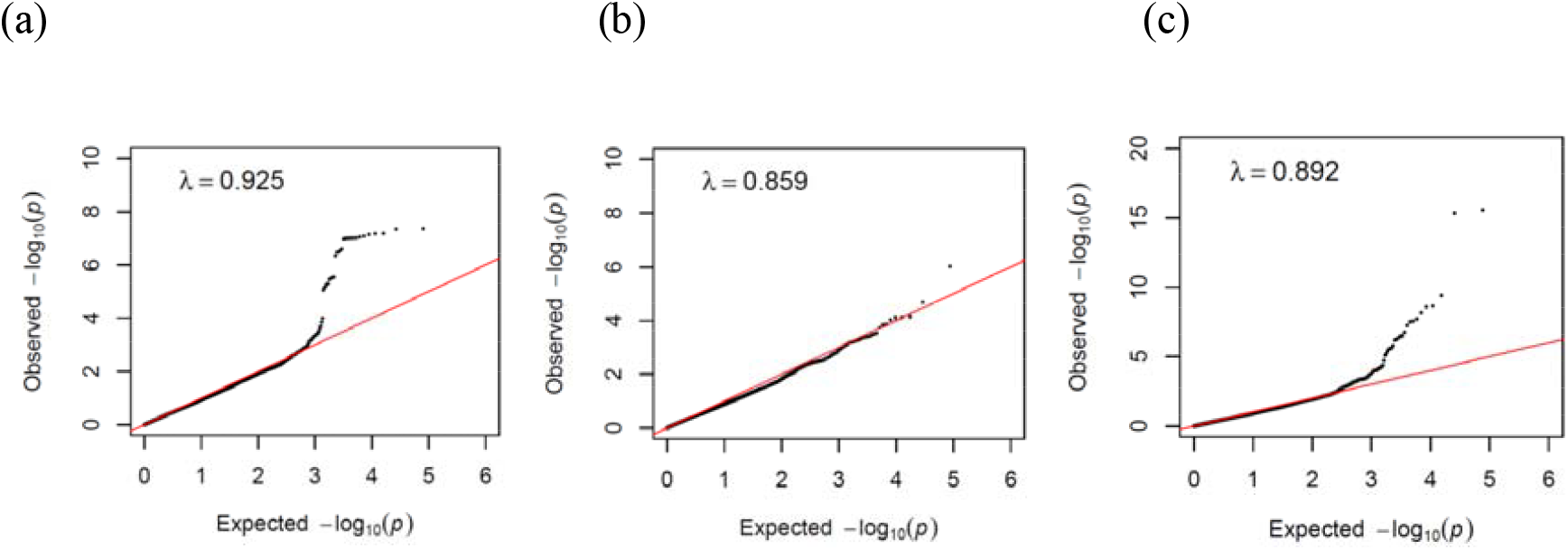
Quantile–quantile plots of the genome-wide association studies for genetic effect of sow on NBA for (a) Line A (b) Line B (c) Line C.

Manhattan plots for genetic effect of sow on NBA for the three lines are shown in Figure 2.

Significant peaks were identified for lines A, B, and C. For line A, 18 significant SNPs were located on chromosome 6, spanning positions near 151.23Mb with -log10(P-values) ranging from 6.34 to 7.37. For line B, one significant SNP was identified on chromosome 1 at position 192.08Mb with a -log10(P-value) of 6.03. For line C, 16 significant SNPs were detected, with -log10(P-values) ranging from 6.21 to 15.56. These SNPs were located on chromosome 1 (positions 14.34Mb to 14.45Mb), chromosome 2 (position 150.48Mb), chromosome 3 (positions 93.80Mb to 94.37Mb), chromosome 5 (position 83.05Mb), and chromosome 9 (positions 117.86Mb to 137.15Mb). The λ of the QQ plot regression were 0.925, 0.859, 0.892 for line A, B, and C, respectively, indicating slight deflation for all three lines.

## Discussion

Over the past few decades, several studies have investigated the influence of paternal components, both genetic and non-genetic, on sow reproductive performance. In earlier research, See et al. [1]reported that although the service sire effect was small, it was significant, accounting for 1 to 2% of the total variation in number born alive in Landrace, Hampshire, and Spotted populations. More recently, Cieleń and Sell-Kubiak [3] reviewed studies assessing the proportion of phenotypic variance in reproductive traits explained by the service sire effect and found that the genetic component accounted for 1 to 2%. The non-genetic service sire effect was slightly higher, ranging from 1% to 5.5%, though this may partly reflect confounding with genetic effects in models where the service sire genetic component was excluded. The heritability estimates for the service sire effect on NBA reported in our study fall within the range of previously published values, indicating low heritability for the service sire effect on litter size in swine.

In our study, we modeled the service sire and sow genetic effects with relationships between them. This approach has not been commonly reported in previous studies, possibly due to software limitation or the complexity of the modeling process. Having service sire as uncorrelated random effect without considering the additive relationship, Wolf and Wolfová [34] reported sow heritability of 0.05 to 0.07 for number of stillborn and number of piglets lost from birth until weaning in Czech Landrace and Czech Large White population. The proportion of variance due to service sire was very low between 1% and 1.2% for number of stillborn and between 0.8 and 1.6% for piglets loss. Chen et al. [4] studied number born alive and weaning performance in U.S. Landrace, Yorkshire, Duroc, and Hampshire populations. For number born alive, heritability estimates for the sow effect ranged from 0.08 to 0.10, while proportion of variance due to service sire effects ranged from 2% to 4%. Serenius et al. [5] investigated the effect of service sire on litter size in Finnish Landrace and Large White populations. They reported the proportions of variance due to service sire effect on litter size were near zero whereas the heritability of sow effect ranged between 0.11 and 0.15. Serenius et al. [6] reported that the proportion of variance due to the service sire effect were 1% to 2% for litter size traits, 1% to 4% for piglet survival traits, and noticeably higher for gestation internal (5% to 9%). Cielen’ et al. [8] investigated litter size variability in Landrace using a model that included both genetic and non-genetic service sire effect, while assuming no relationships between service sire and sow. They also reported the genetic variance attributable to the service sire accounted for less than 2% of the total phenotypic variance.

Considering the additive covariance among the service sire and sow effects, Van der Lende et al. [9] analyzed one purebred sire line and two purebred maternal lines and one F1 line in Netherlands. For purebred genetic parameter estimates, they reported heritability estimates for service sire effect on litter size ranging from 0 to 0.04, whereas sow heritability estimates were nearly threefold higher ranging from 0.07 to 0.12. The genetic correlation between service sire and sow effects ranged from -0.47 to 0.27. Kim et al. [10] reported sow and service sire genetic parameters for litter size and birth weights in Korea Yorkshire population. Estimates of heritability of sow effect for litter size were lower (0.18 to 0.19) compared to birth weight traits (0.25 to 0.39). Service sire heritability estimates ranged from 0.01 to 0.02 for all traits. The genetic correlation between service sire and sow effects were 0.01 to 0.11 for litter size and from -0.15 to 0.54 for birth weights traits.

In our study, the permanent environmental variance accounted for less than 1% of the total phenotypic variance. Several studies reported that permanent environmental effects of the service sire were not included in the models with some studies indicating preliminary analysis showed a low estimate of this effect [4,9,10,34].

The heritability estimates for semen motility (0.10 to 0.12) and abnormal morphology (0.17 to 0.28) in this study fall within the range of values reported in the literature. Morphology can be analyzed based on individual traits or as a composite measure. The genetic parameter estimates summarized herein focus on morphology defined as a total score.

For Duroc, Gruhot et al. [11] studied about 400 boars and found heritability estimates ranged from 0.08 to 0.24 for semen traits using data collected during the summer season. Ogawa et al. [12] analyzed semen traits from 900 boars with 46K semen collection records. The heritability was about 0.20 with repeatability of 0.46 for proportion of morphologically normal sperm. For Landrace and Large White, Wolf [7] estimated heritability in Czech Landrace and Large White pigs. They reported heritability for motility and morphology were around 0.10 based on 37K to 51K ejaculates from about 800 boars per breed. Marques et al. [35]reported heritability of 0.18 for motility and 0.20 for morphology. Krupa et al. [36] using 57K sperm collections records from 1,209 Czech Landrace and Large White boars and reported heritability of 0.11 to 0.14 for motility and 0.22 to 0.24 for morphology. Sa et al. [13] analyzed semen traits collected from 450K ejaculates on about 5.7K boars. For motility, estimates of heritability ranged from 0.20 to 0.24 with repeatability ranged from 0.35 to 0.58. For total morphological abnormalities, the estimate of heritability was 0.21 with repeatability of 0.52. For other sperm morphology traits, heritability ranged from 0.11 to 0.27 with repeatability ranged from 0.20 to 0.65. Overall, the reported heritability estimates for morphology from literature and this study were similar or higher than estimates for motility across breeds.

The negative genetic correlations between motility and morphology were observed in the current study with a range from -0.64 to -0.34. The negative genetic correlations were also reported in literature [7,11,35,36]. For example, the genetic correlation of -0.66 was reported in Marques et al. and estimates between -0.42 and -0.35 were reported in Krupa et al. [36].

For genetic correlations between litter size and semen traits, Wolf [7] concluded that estimates could be breed-specific. Ogawa et al. [12] found the genetic correlation between normal morphology and litter size was low with high SE but in favorable direction. Those correlations were 0.12, 0.18 and -0.17 for total number born, born alive, and stillborn, respectively. In our study, estimates of GEBVs correlation between litter size and semen quality were low, but overall motility positively affected litter size while abnormal morphology negatively affected litter size.

In the current study, there was only one significant marker associated with NBA for service sire effect in line B. However, the -log10(P-Value) was close to the significance threshold. Although there were more significant markers associated with NBA for sow effect, the objective of our GWAS analysis was to focus on the service sire effect. To our knowledge, this study was the first one investigating service sire effects for litter size applying ssGWAS with APY approach in pig dataset.

## Conclusions

Based on large data sets of 1M ejaculates from 50K boars from three terminal sire lines, this study reports low to moderate heritability and moderate to high repeatability estimates for semen motility and total abnormal morphology traits. Moderate and favorable genetic correlations between semen quality traits indicate that selection for semen motility might be favorable for improving morphology. To our knowledge, this study is the first to investigate service sire effects on litter size in pigs using genomic data through ssGWAS with the APY approach. The genetic impact of the service sire effect on litter size appeared to be small, as indicated by the low heritability estimates and the limited number of significant markers detected. Although paternal genetic contributions to litter size were small compared to maternal genetic contributions, selecting on service sire effects on litter size in addition to semen quality traits will improve overall reproductive success.

## Authors’ contributions

CYC and JH conceived and designed the study. CYC performed the analyses and wrote the first draft. CYC, MK, and AB prepared the data set, DL provided analysis support and software. CYC, MK, AB, DL, and JH provided comments and revised the manuscript. All authors read and approved the submitted version.

## Declarations

### Ethics approval and consent to participate

Not applicable

### Consent for publication

Not applicable

### Availability of data and materials

The dataset used in this study was obtained from a pre-existing dataset owned by Genus PIC and not publicly available.

